# Conditioning electrical stimulation enhances functional rewiring in a mouse model of nerve transfer to treat chronic spinal cord injury

**DOI:** 10.1101/2025.01.17.633666

**Authors:** Juan Sebastián Jara, Marwa Soliman, Amanda Bernstein, Paola di Grazia, Adam R. Ferguson, Justin M. Brown, Abel Torres-Espín, Edmund R. Hollis

## Abstract

Nerve transfer surgery is a state-of-the-art surgical approach to restore hand and arm function in individuals living with tetraplegia, significantly impacting daily life. While nearly a third of all individuals with chronic SCI may benefit from this intervention, variability in outcomes can limit the functional impact. A bedside-to-bench approach was taken to address the variable response of tetraplegic individuals to nerve transfer surgery. We used a hierarchical multiple factor analysis to evaluate the effects of conditioning electrical stimulation (CES) on outcomes in a mouse model of nerve transfer to treat chronic cervical spinal cord injury. We found that CES of donor nerves one week prior to nerve transfer surgery enhanced anatomical and functional measures of innervation of targeted muscles. Furthermore, CES increased the rate of recovery of naturalistic behavior. While the model has some limitations due to the small size of the rodent, our results support the use of CES as an effective approach to improve outcomes in clinical nerve repair settings.

## Introduction

The translation of scientific discoveries from the laboratory to the clinic is a complex and multifaceted process during which most strategies fail. Bridging the gap between basic and translational neuroscience studies and their application in clinical settings requires a dynamic and iterative approach. A bedside to bench and back again approach (Hampton, 2017), characterized by a cyclical information flow between the clinic and the laboratory, holds immense potential for addressing the clinical challenges and meeting the needs of individuals living with chronic neurological injury. This iterative process allows for the refinement and optimization of experimental models and therapeutic interventions based on clinical observations, while also generating new insights and discoveries that can inform and guide clinical practice. In this vein, we have taken an interdisciplinary approach to test the therapeutic effect of conditioning electrical stimulation (CES) in a novel mouse model of nerve transfer surgery to treat chronic spinal cord injury (SCI). Nerve transfer is a current surgical intervention for individuals living with chronic tetraplegia that involves the transection and coaptation of expendable, peripheral donor nerves arising above the level of injury with recipient nerves from at or below level. This approach works through peripheral axon regeneration and subsequent re-innervation of muscles that had lost supraspinal motor input (Brown, 2011; Fox, 2015; Bertelli, 2017).

While nerve transfer surgery has shown success in treating cervical SCI, there is still significant variability in the recovery of dexterity among recipients (Khalifeh, 2019). This variability can be attributed, in part, to the level and extent of individual injuries. However, the rate and extent of axon regeneration play crucial roles in determining clinical outcomes. Numerous studies have shown that the rate and extent of regeneration in peripheral nerves can be enhanced through a conditioning injury (McQuarrie, 1973), which induces a distinct transcriptional program promoting robust neuronal and non-neuronal responses (Tsujino, 2000; Costigan, 2002; Boeshore, 2004; Seijffers, 2007; Stam, 2007; Chandran, 2016).

Electrical stimulation has emerged as a potential adjunct therapy to enhance nerve regeneration. Low-frequency stimulation delivered directly to intact peripheral nerves has been shown to upregulate the expression of regeneration-associated genes (RAGs) associated with conditioning injury (English, 2007; Geremia, 2007; Senger, 2018; Senger, 2019). Notably, pre-injury CES has been found to be more effective in eliciting regeneration compared to post-injury stimulation (Senger, 2020). Furthermore, clinical trials have demonstrated the safety of direct electrical stimulation in reparative surgeries involving median or digital nerves (Gordon, 2010; Wong, 2015).

Considering the limitations and variability observed in the current surgical intervention for SCI, we developed a translational model of nerve transfer surgery in mice to evaluate the effects of CES on functional outcomes. This model was used to investigate the potential of pre-surgical electrical stimulation to enhance axon regeneration, functional reinnervation, and behavioral recovery. Using a hierarchical multiple factor analysis (HMFA), we observed that CES reduced the variability in outcomes in part due to greater functional connectivity and a faster recovery of naturalistic movements. By modifying clinical techniques at the bench before translating back to the bedside, our results support the use of CES to improve the efficacy and outcomes of nerve transfer surgery for the treatment of chronic tetraplegia.

## Materials and methods

### Animals

All animal experiments were approved by the Weill Cornell Medicine Institutional Animal Care and Use Committee. C57BL/6J mice (The Jackson Laboratory) were housed under 12-hour light/dark cycle, humidity 39-48%, and average temperature of 21.7°C with food and water *ad libitum*. For histological assay of sensory axon regeneration, transgenic mice (C57BL/6J background) expressing Cre-recombinase under the control of parvalbumin promotor (*Pvalbtm1(Cre)Arbr*) were crossed with transgenic *Ai14-LSL-tdTomato* mice (The Jackson Laboratory) to generate *Pvalb::tdTomato* mice with genetically labeled, large-diameter sensory neurons.

### Experimental overview

Prior to SCI C57BL/6J mice were food restricted to 80–90% of their free-feeding bodyweight, then trained on the single pellet reach task and tested for baseline performance on other behavioral tests as indicated below. Dominant forelimb was identified for single pellet reach and surgical procedures targeted this forelimb. After training on the forelimb reach and recording baseline assessment on grip strength, adhesive removal, and pasta handling, mice underwent SCI (described below). Twelve weeks after SCI, chronically injured mice underwent behavioral testing followed by random allocation to CES or sham stimulation. One week after CES or sham stimulation, mice underwent unilateral musculocutaneous to median nerve transfer surgery. Functional recovery was evaluated beginning 4 weeks after nerve transfer with twice weekly training on single pellet reach over the next 16 weeks. Grip strength, adhesive removal, and pasta handling tests were performed at 4, 8, 12, and 16-weeks post-nerve transfer. Upon completion of rehabilitation, mice underwent electromyography (EMG) recordings and retrograde tracing (described below).

### Spinal cord injury

Twenty adult C57BL/6J mice (8-10 weeks old, both sexes) were deeply anesthetized with isoflurane, the skin was shaved and cleaned, an injection of bupivacaine (0.25% solution) was made prior to incision, and a laminectomy of cervical 6 vertebrae was performed to expose cervical 7 spinal level (C7). A dorsal over-quadrantsection (DoQx) micro-scissor lesion at C7 was used to transect the dorsal columns bilaterally and the dorsolateral spinal cord unilaterally at a depth of 1 mm from the dorsal surface of the cord (Figure 1a). This injury is restricted to a single spinal level, targets the corticospinal and rubrospinal descending motor pathways, and results in a permanent deficit in dexterous prehension. After surgery the overlapping spinal muscle and fascia were re-apposed using sterile absorbable suture. The skin was closed with sterilized wound clips and mice were given the analgesic buprenorphine (0.1 mg/kg) subcutaneously at the time of surgery and twice a day for 3 days.

**Figure 1.**
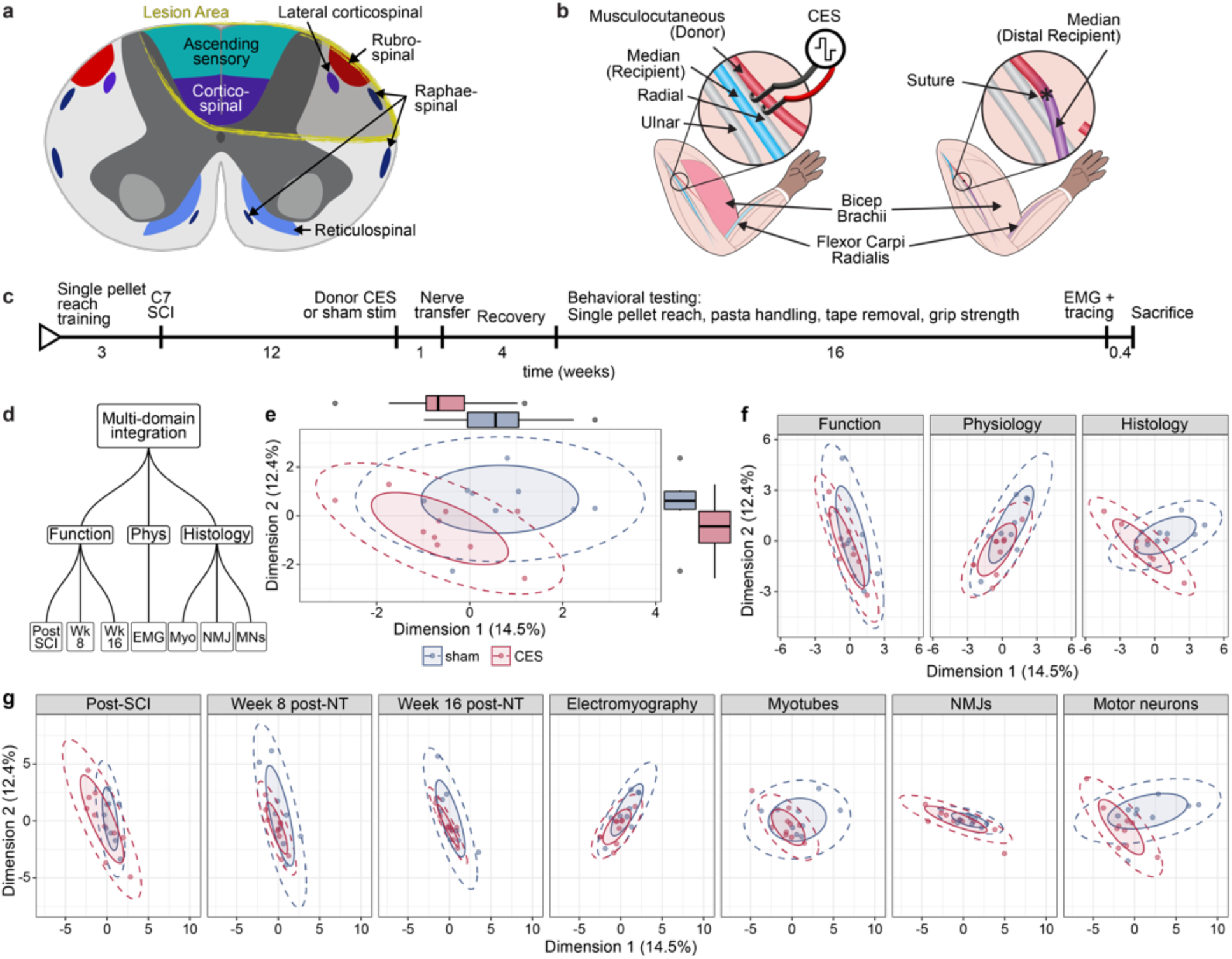
CES alters outcomes in a mouse model of nerve transfer surgery to treat chronic SCI. **a)** Dorsal over-quandrantsection transects descending corticospinal and rubrospinal tracts as well as ascending, dorsal column sensory axons. **b)** Schematic of donor nerve CES in a mouse musculocutaneous to median nerve transfer model. **c)** Timeline of experiment using nerve transfer to treat chronic SCI in mice. **d-g)** Multi-domain integrative analysis across function, physiology, and histology parameters **d)** demonstrate overall changes between sham and CES (p = 0.019) **e)**. These are also observable at the level of each domain in **f)** and at the level of each group of variables in **g)**.

### Conditioning electrical stimulation

Twelve weeks after spinal cord injury, mice were randomly assigned to CES or sham stimulation. Mice were deeply anesthetized with isoflurane, the dominant forelimb was shaved and cleaned, bupivacaine was injected over the pectoralis muscle, an incision was made, and muscles were retracted to expose the terminal branches of the brachial plexus. A pair of stainless-steel hook electrodes (Microprobes) were placed over the musculocutaneous nerve and 20 Hz, 0.2 ms pulse low threshold direct electrical stimulation was performed for 1 hr with current set to twice the motor threshold. Sham stimulation involved hook electrode placement for 1 hr with no electrical stimulation. During this procedure one mouse did not survive anesthesia induction.

### Nerve transfer surgery

One week after CES or sham stimulation, mice underwent musculocutaneous to median nerve transfer in the dominant forelimb (Figure 1b). Brachial plexus branches were exposed as above, then median and musculocutaneous nerves were transected distal to *biceps brachii* innervation and proximal musculocutaneous was coapted to distal median nerve using 10-0 nylon suture (Ethilon 2810G). The proximal median nerve stump was sutured to adjacent musculature to prevent regeneration.

### Single pellet reach behavior

Single pellet reach behavior was conducted to assess dexterous prehension in mice. The acrylic behavior box had three slots (7 mm wide) positioned on the left, center, and right sides of the front wall. Flavored food pellets weighing 20 mg (Bioserv, F05301) were placed at 12 mm from the inside wall of the box on a platform level with the box floor. The dominant forelimb was determined during the first training session. Mice were trained over 15 training sessions spread across 3 weeks, each session consisting of 25 trials. Eleven weeks after SCI, mice were tested twice prior to nerve transfer and then weekly for 16 weeks post-transfer. Only trials with successful pellet contact were counted, and the success rate was calculated as the percentage of trials with successful pellet retrieval and eating. High-speed video of the task was recorded on a Basler Ace camera (acA1440-220um) with a 12 mm lens at 327 fps, 720 x 540 px resolution for quantification of forelimb reach trajectories using the markerless pose estimation AI DeepLabCut (Mathis, 2018). In DeepLabCut (version 2.2b7), seventeen videos were used to compose the training set. From each video, at least 70 frames were extracted where digits 2-5, the nose, and the pellet were labeled. These frames were used to refine the pre-trained network (ResNet-50) to predict features of interest in unseen videos and generate x- and y-coordinates for each label throughout the video. After training the DeepLabCut neural network, locations of forelimb digits, paw, and pellet were marked in recordings of each reach trial. The raw x and y spatial position for markers in each frame was extracted, pre-processed, and quantified using a kinematic deviation index (KDI) we describe elsewhere (Torres-Espin, 2024). In brief, KDI is a unitless summary metric derived using a machine learning workflow denoting global changes in the 3D trajectory of all markers with respect to pre-injury baseline performance. A KDI value is extracted for each trial and each animal and used as outcome measure of single pellet reach performance.

### Adhesive removal task

Adhesive tape (3x4 mm) was stuck on the forepaw before gently placing the mouse in an acrylic cylinder (13 cm diameter, 18 cm height). Each animal was tested bilaterally and the time taken to sense the tape (paw shake, sensory component) and to completely remove it (motor component) were recorded (Bouet, 2009).

### Grip strength

A grip strength meter (Columbus Instruments) was used to evaluate neuromuscular function after nerve transfer. Mice were tested by being gently lifted to grasp the pull bar assemblies (inclined 45°) attached to the force measurement device, and slowly pulled away in a horizontal direction. Average grasp force was calculated from 3 trials per animal per session.

### Pasta handling

Animals were placed in an 8 x 6.9 cm box with an angled mirror below and filmed for 20 minutes. Videos were recorded using a Basler Ace camera (acA1440-220um) with a 12 mm lens at 227 frames per second. DeepLabCut (version 2.2b7) was used to track pasta and mouse body parts during the 20 minutes to generate quantitative measures of impairment such as the angle of the pasta and the distance between the paws. In DeepLabCut, 61 videos were used to compose the training set. From each video, at least 30 frames were extracted where body parts in both the front and mirrored view were labeled. The angle of the pasta was determined by calculating the angle between the pasta and a straight line from the nose to the midpoint of the hind paw base of support in the mirrored view using Python. The density distribution for angles in the injured paw and the non-injured paw were estimated using a Gaussian kernel with a bandwidth of 2 angles. All angles were oriented so that those from the injured side were positive angles (0-180°) and from the uninjured side were negative angles (- 180°-0).

### EMG recordings

Electrophysiological recordings using needle electrodes in the forelimb (Pollari, 2018) were used to evaluate functional re-innervation after nerve transfer. Mice were anesthetized with ketamine/xylazine cocktail (100mg/kg and 10mg/kg respectively). Two electrodes (27-gauge needle) were placed subcutaneously over site of the nerve transfer surgery to stimulate the coapted nerve at mid-humerus level with 0.1 Hz and 0.1 ms pulse duration. Two recording electrodes (29G needle, AD Instruments # MLA1213) were located subcutaneously over the *flexor carpi radialis* muscle (FCR) at mid forearm level, and a reference electrode was placed in the walking pad of the same recorded forelimb (3 mm depth approximately). Recordings were performed using a differential AC amplifier (A-M Systems) and PowerLab 8/35 data acquisition system, LabChart was used for visualization. Control EMG recordings for each mouse were recorded from stimulation of contralateral, intact median nerve. Rectified EMG signals were averaged (6 recordings per stimulation intensity) to determine motor evoked potential (MEP) thresholds. Responses were recorded starting from the minimum intensity required to elicit an M-wave followed with increasing intensities until the maximum M-wave amplitude was achieved (threshold).

### Tracing

Following EMG recordings, the retrograde and transganglionic tracer cholera toxin B subunit (CTB, 1% wt/vol in dH_2_O) was injected into the median nerve distal to the nerve coaptation site (or into contralateral intact median nerve). Mice were transcardially perfused 3 days later and tissues were isolated for histology.

### Axon regeneration assay

Six adult *Pvalb::tdTomato* mice (8–10-week-old, both sexes, littermate controls) underwent CES or sham stimulation of musculocutaneous nerve as above, followed one week later with musculocutaneous to median nerve transfer. Mice were transcardially perfused 7 days after nerve transfer and coapted nerves were isolated for histology. To evaluate axonal regeneration, td-tomato-positive fibers were identified in confocal images and counted every 250µm from the nerve transfer site to the distal end of the recipient nerve.

### Tissue processing

Fixed tissue was cryoprotected with 30% sucrose in PBS and then cut into 20 µm-thick horizontal sections that were blocked in phosphate buffer (PB) containing 5% Donkey serum and 0.2% Triton X-100 for 60 minutes at room temperature. Tissue sections were incubated overnight at room temperature with primary antibodies in PBS and 5% Donkey serum and 0.2% Triton X-100. Sections were then incubated with appropriate Alexa Fluor (ThermoFisher) secondary antibodies (1:500 dilution) in PBS for 2 hours at room temperature. Sections were counterstained with DAPI and mounted with Fluoromount-G (ThermoFisher Scientific). The following primary antibodies and dilutions were used: mouse anti-NeuN (1:200, Millipore # MAB377)), rabbit anti-ATF-3 (1:200, Novus #NBP1-8581), mouse anti-laminin (1:50, Sigma-Aldrich #CC095-M), goat anti-CTB (1:200, List Biological Laboratories, #703), rabbit anti-DsRED (1:200, Takara # 632496), rabbit anti-Synaptophysin Sp11 (Thermo Fisher Scientific # MA5-14532), mouse anti-Myosin heavy chain type IIB (MHC IIb, 1:100, Developmental Studies Hybridoma Bank, #BF-F3), Alexa Fluor 488 conjugated α-Bungarotoxin (1:500, Thermo Fisher Scientific #B13422).

### Myotube analysis

Transverse sections of *FCR* stained with a combination of anti-laminin and anti-MHC IIb were processed in FIJI. Myotube outlines in thresholded images were used to generate masks of individual fibers for cross-sectional area analysis.

### Statistical analysis

Multi-domain integration analysis was performed using hierarchical multiple factor analysis (HMFA) (Pag*ès*, 2014) as implemented in the FactorMineR R package (Lê, 2008). Data were organized in three hierarchical levels, the global multi-domain integration level, 3 subsequent levels containing function, electrophysiology, and histology data separately, and a bottom level that specifies the considered data tables. For function, we analyzed chronic, 12-week post-SCI, and then 8 weeks and 16 weeks after nerve transfer as separate tables including grip strength, the pasta drops, pasta paw preference, tape removal time, tape detection time, KDI, and summary statistics of the paste angle distribution (minimum, maximum, first, second and third quartiles, kurtosis, skewness and standard deviation). For electrophysiology, we analyzed EMG parameters including area under the curve, and threshold. For histology, we included myotubes and neuromuscular junction density summary statistics (minimum, maximum, first, second and third quartiles, kurtosis, skewness and standard deviation), and motor neuron counts in a 2D grid. Previous HMFA, missing data (9.9% of the total data), was imputed using the *imputeMFA()* function in the missMDA R package (Josse, 2016), which performs imputation by a regularized iterative MFA algorithm. The first 3 dimensions had an eigenvalue > 1, amounting to 38% of the total variance. To perform inference and test on the hypothesis of differences between groups in the scores first plain of the global and partial solution (components 1 and 2) we performed a multivariate analysis of variance (MANOVA) with Wilks test. Univariate hypothesis test for the difference between groups in each component was conducted using Wilcox sum of rank test.

To perform inference on the hypothesis that CES had an effect on the different metrics compared to sham stimulation, distinct statistical methods were used. For KDI over time, a linear mixed model (LMM, using the lme4 and lmerTest R packages (Bates, 2015; Kuznetsova, 2017)) with treatment Group (CES, sham), Timepoint of measurement, and the interaction of Group X Timepoint as fixed terms, and subject random term with random intercept and slope over Timepoint. An F-test with Satterthwaite’s approximation of degrees of freedom was conducted to test the significance of each fixed term. For all metrics that are continuous curves (e.g., a density distribution of MN over rostrocaudal distance), we used generalized additive models (GAM). These allow for considering two shortcomings of LMMs and ANOVA methods when used for continuous curves: the assumption of a linear association, and the assumption of normality. For the density angles in the pasta test, a mixed effect Beta regression GAM was fitted for each Timepoint and paw (injured, non-injured) comparing the Groups, the Angle X Group interaction, and specifying subjects as random effect, and a smoother spline in angle.

For the normalized amplitude of the EMG response, a mixed effects Beta regression GAM smoothed over threshold with a 4-order spline were performed as above comparing Groups. For the density of volume and intensity colocalization in the neuromuscular junction analysis and the density of the myofiber area, mixed effects Gaussian regression GAMs and F-test were performed as above comparing Groups. The relative counts of axons (to the reference point of 0.5 mm proximal to the coaptation point) in the axonal regeneration assay were modeled as a mixed effects Beta regression with a 4-order smooth spline over the regeneration distance starting at 0.25 mm distance distal to the coaptation point. For the count density of motor neurons over the rostrocaudal axis, a mixed effects Beta regression GAM was fitted. In all GAMs, a t-test was used to test the significance between Groups, and a Chi squared deviance test comparing a model with and without the interaction was used to assess significance of interactions. Injured and non-injured paws were analyzed separately. In all analysis, a p-value < 0.05 was considered significant. All density and GAM plots are represented as the average smooth ± 1 standard deviation. GAMs were fitted using than *bam()* function the mgcv R package (Wood, 2017).

For electrophysiology, we analyzed the ratio of the ipsilateral to contralateral sides of EMG parameters including area under the curve, latency, and threshold. For each of these parameters, unpaired T-tests were run (GraphPad Prism 10.3.1) to test the significance between the control and treatment groups. A 2-way ANOVA with Bonferroni’s multiple comparisons test was run to test the significance between the control and treatment groups on total time to tape removal.

## Results

### CES improves outcomes after nerve transfer surgery to treat chronic cervical SCI in mice

Using a holistic Hierarchical Multiple Factor Analysis (HMFA) integrative approach to evaluate outcomes across modalities (functional, physiological, and histological) in the several collected data at once, we found that CES significantly alters outcomes globally (MANOVA Wilks lambda = 0.61; F _(2, 16)_ = 5.07, p-value = 0.019) and trends towards reduced outcome variability after nerve transfer (Figure 1d-g). In chronic SCI, prior to CES, there is no difference in function, yet at 8- and 16-weeks post-nerve transfer, CES results in less variability in behavioral outcomes in our mouse model and a significant change in global behavioral function (MANOVA Wilks lambda = 0.64; F _(2, 16)_ = 4.39, p-value = 0.03) (Figure 1f). Physiological and histological measures taken after 16 weeks of recovery also show differences from control, sham stimulation, reaching statistical significance for histology (MANOVA Wilks lambda = 0.63; F _(2, 16)_ = 4.514, p-value = 0.028) (Figure 1f).

### CES enhances peripheral regeneration after nerve transfer surgery in mice

To confirm that CES enhances axon regeneration in a mouse model of nerve transfer, we evaluated the regeneration of large diameter, parvalbumin expressing sensory axons acutely after musculocutaneous to median nerve transfer using *Pvalb::tdTomato* transgenic mice (Figure 2a-c). Musculocutaneous donor nerve was stimulated 1 week before nerve transfer, during which musculocutaneous and median nerves were transected distal to *biceps brachii* innervation and proximal musculocutaneous was coapted to the distal median nerve. We evaluated sensory axon regeneration 1 week after nerve transfer and found a robust enhancement of large-diameter, tdTomato^+^ sensory axon regeneration in CES mice (Group effect GAM p-value = 0.0168) (Figure 2d).

**Figure 2.**
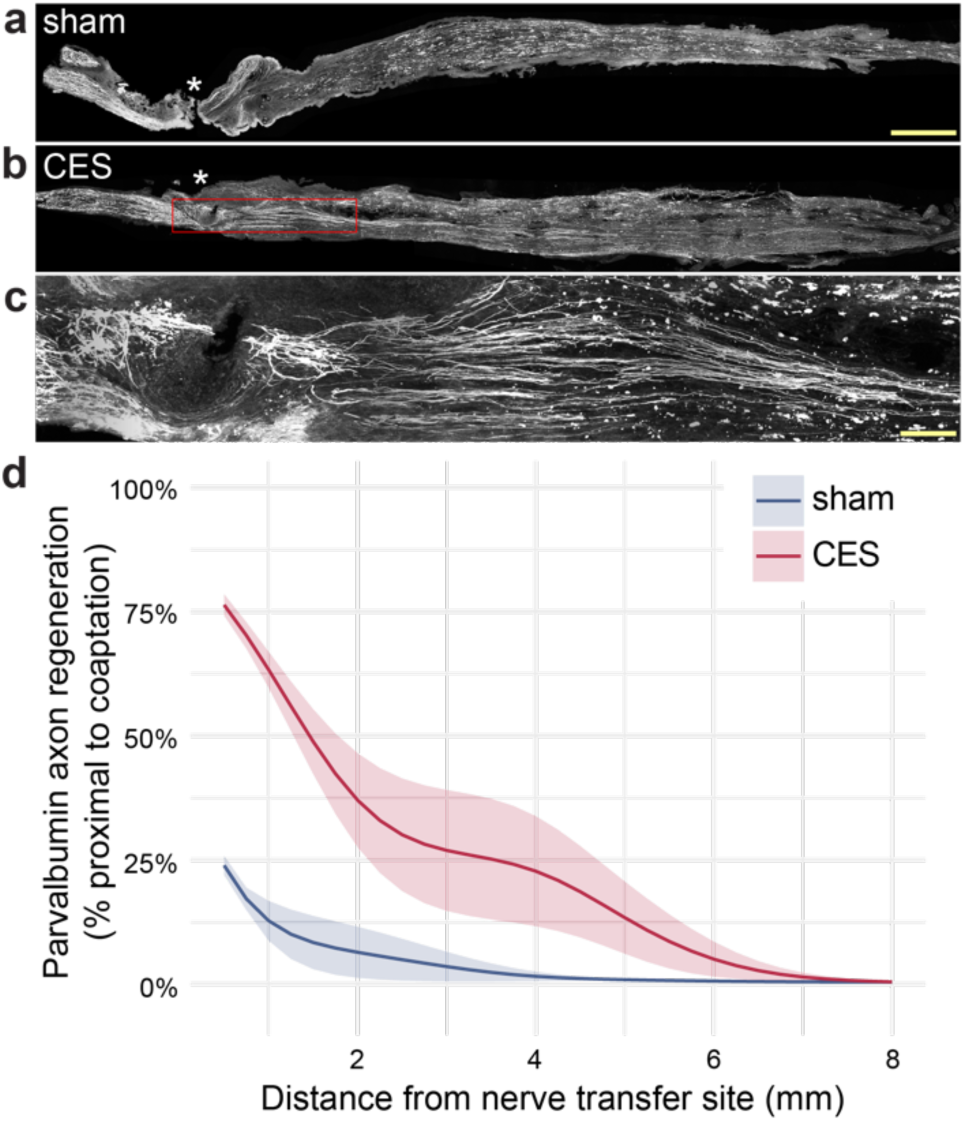
CES enhances regeneration in a mouse model of nerve transfer surgery. CES significantly enhances regeneration of parvalbumin expressing sensory neurons at 7 days post-transfer (mean ± sem). Scale bars = 100 um (a,b,f), 20 um (c), 500 um (d,e). Group effect GAM p-value = 0.0168; Group X Distance interaction GAM p-value < 0.01. Data presented as mean ± sem.

### Limits of CES in a mouse model of nerve transfer to treat chronic SCI

To evaluate the functional effects of CES on outcomes in our mouse model of nerve transfer to treat chronic tetraplegia, we performed CES or sham stimulation in mice with chronic C7 dorsal over-quadrantsection (DoQx) SCI. This injury interrupts the main corticospinal tract bilaterally as well as the dorsolateral corticospinal tract and rubrospinal tract unilaterally. Prior to SCI, mice were trained on single pellet reach behavior, then tested on a battery of assessments before undergoing DoQx lesion to target the dominant forepaw used for pellet reach. Impairment of forelimb function was tested at 7 weeks post-SCI. After 12 weeks, mice underwent CES or sham stimulation followed one week later by musculocutaneous to median nerve transfer, sacrificing musculocutaneous function. After 4 weeks of recovery, mice began twice weekly testing on forelimb reach and impairment was tested on adhesive removal task, grip strength, and pasta handling.

Over 16 weeks of twice weekly training, mice showed no recovery of success on the single pellet reach task (Figure 3b). High speed video analysis was used to generate the kinematic deviation index (KDI). KDI was used to evaluate the similarity of reach trajectories to successful pellet retrieval by the same animal prior to SCI. No differences in KDI were observed over the course of recovery (Figure 3c). Additionally, we tested mouse grip strength over time and found no differences between groups (Figure 4e). Furthermore, musculocutaneous to median nerve transfer impaired overall grip strength, which depends on innervation of musculocutaneous upper arm targets.

**Figure 3.**
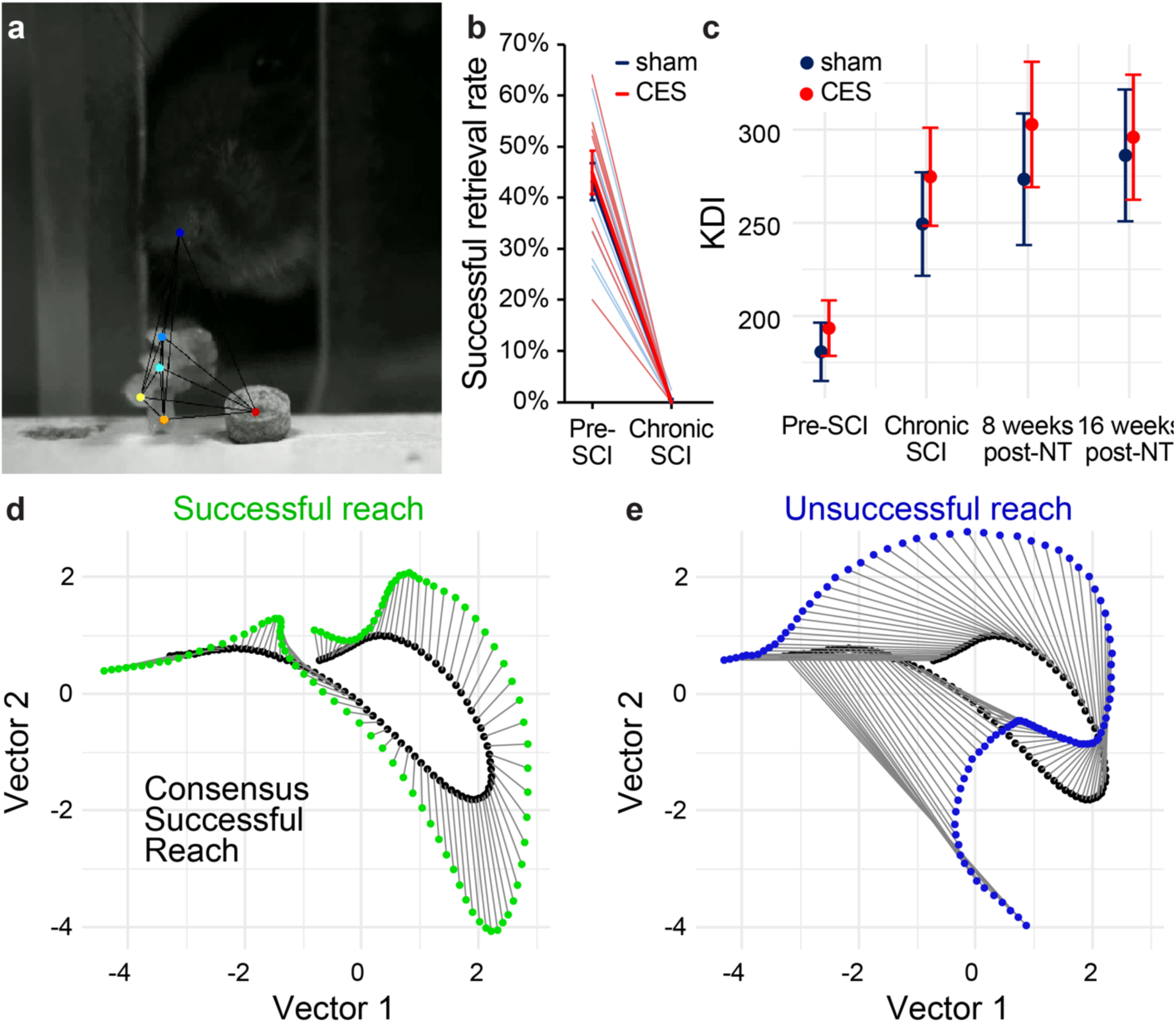
C7 DoQx SCI chronically impairs precision forelimb prehension. **a)** Mice were trained and tested on single pellet reach and digits were labeled and tracked with DeepLabCut. **b)** Successful reach was completely abolished on single pellet reach in chronically injured mice after C7 DoQx SCI. **c)** KDI values generated from the Euclidian distance for each trace sample between the test trial and the pre-SCI consensus on the 3 PCs showed no effect of CES on reach trajectory. **d,e)** Examples of successful and unsuccessful individual reach trials compared to pre-SCI consensus successful reaches for one animal. Data presented as mean ± sem.

**Figure 4.**
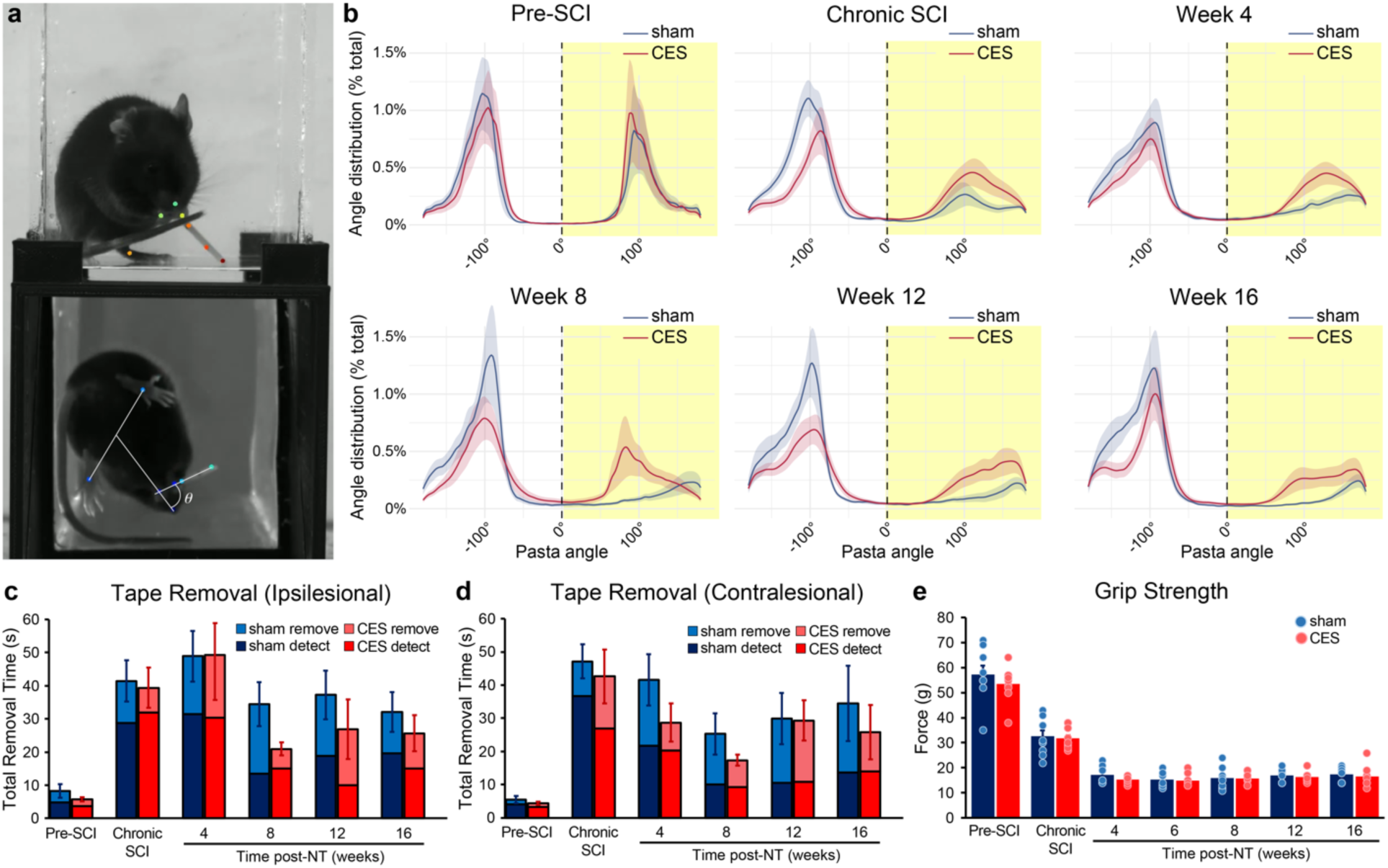
CES enhanced the rate of recovery of naturalistic movements. By 8 weeks after nerve transfer, CES treated mice showed more rapid recovery of **a,b)** pasta handling (Group X Angle interaction GAM p-value < 0.01 in the transfer side) and **c)** tape removal from the ipsilesional, nerve transfer paw, with limited effects on contralesional, intact median nerve paw tape removal **(d)**. **e)** Musculocutaneous to median nerve transfer severely impacted forelimb grip strength. Data presented as mean ± sem.

### CES enhances recovery of naturalistic movements after nerve transfer

To determine effects of CES on recovery of naturalistic forelimb dexterity after nerve transfer, we used pasta handling and tape removal tasks. Mice showed greater recovery on pasta handling with the affected forelimb by 4 weeks post-nerve transfer with CES than sham stimulation (Group effect GAM: 4 weeks p-value = 0.03; 8 weeks p-value = 0.024, 12 weeks p-value = 0.025; 16 weeks p-value < 0.01) (Figure 4b).

### CES enhances reinnervation of median nerve targets after nerve transfer

To assess the extent of anatomical reorganization after musculocutaneous to median nerve transfer surgery, we characterized innervation and myotube structure in the median nerve target *FCR*. Using immunostaining and 3D reconstructions we observed that CES enhanced the reinnervation of *FCR*. CES resulted in significant recovery of pre-synaptic innervation, with fewer small, fragmented neuromuscular junctions than in sham stimulated animals (Figure 5c). Additionally, unlike in sham-stimulated mice, pre-synaptic synaptophysin labeling was similar to that in the contralateral *FCR* with intact median nerve innervation in CES treated mice (Figure 5e). Transverse sections of *FCR* showed that nerve transfer resulted in a higher density of larger myotubes in CES than sham. This effect is reversed in the contralateral limb with intact median nerve innervation where sham presented with higher density of larger myotubes than CES (Figure 5i,j) (Group X Area interaction GAM for intact p-value < 0.01, for transfer p-value < 0.01).

**Figure 5.**
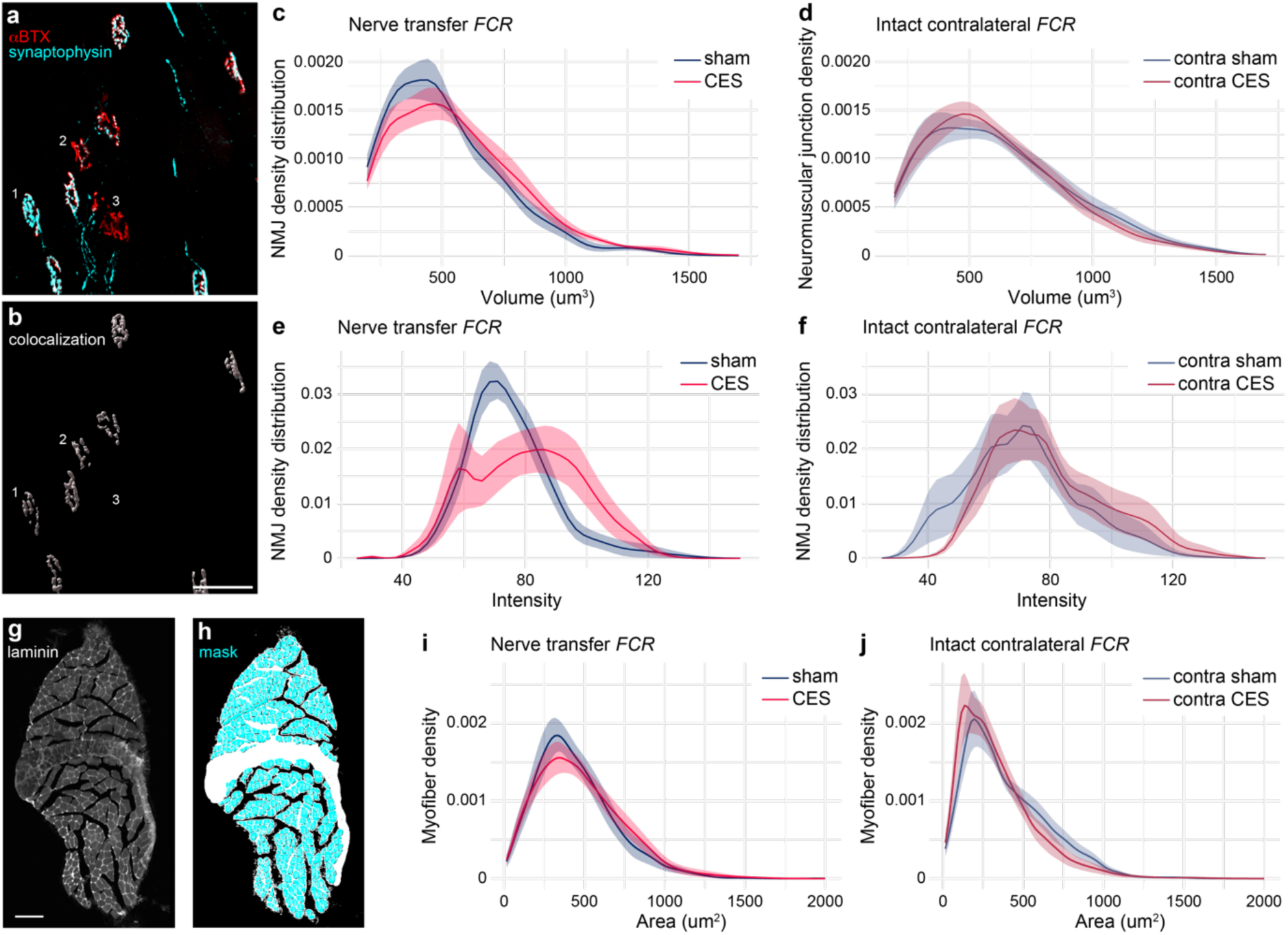
CES enhances neuromuscular junction innervation. **a)** 3D reconstruction of motor endplates labeled shows examples of fully innervated (1), partially innervated (2), and unoccupied neuromuscular junctions (3). **b)** Colocalization of pre-synaptic synaptophysin and nicotinic acetylcholine receptors labeled with α-bungarotoxin (αBTX). **c,d)** CES reduced the number of small motor endplates with a volumetric distribution similar to contralateral intact *FCR*. **e,f)** CES resulted in greater intensity of presynaptic labeling than sham stimulation (Group X Intensity interaction GAM p < 0.01 in **e**, p = 0.076 in **f**). **g,h)** Laminin immunostaining was used to mask individual myotubes for area quantification. **i,j)** Distribution of myotube areas in *FCR* (Group X Area interaction GAM for transfer side p-value < 0.01, for intact site p-value < 0.01). Scale bars = 50 um (b) and 100 um (g). Data presented as mean ± sem.

### Nerve transfer alters the distribution of motor neuron populations innervating the nerve transfer

We used retrograde tracing with CTB to determine the distribution of motor neurons projecting to the transferred nerve. As expected, musculocutaneous to median nerve transfer drives a rostral shift in the population of labeled motor neurons (Site [transfer vs. Intact] X Distance interaction GAM for sham group p-value < 0.01; for CES group p = 0.024), in comparison to the population of motor neurons labeled by tracer injection to the contralateral, intact median nerve. CES alters the overall distribution of motor neurons that project through the nerve transfer (Group X Distance interaction GAM p-value < 0.01) (Figure 6).

**Figure 6.**
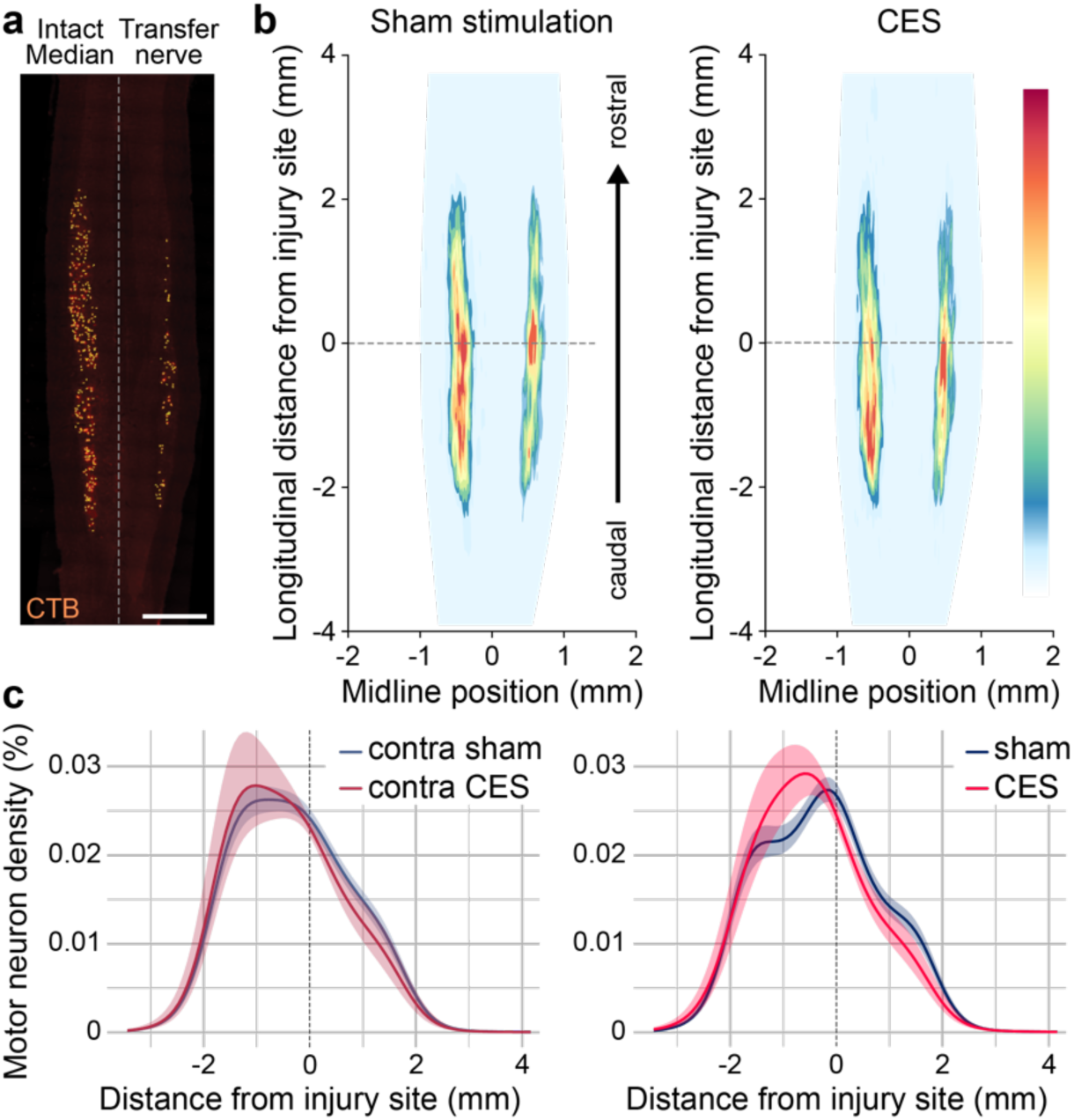
Motor neuron distribution after nerve transfer is altered by CES. **a)** Example of CTB traced motor neurons following injection to musculocutaneous-median nerve transfer or contralateral intact median nerve. **b)** Heat maps show average distribution of all labeled motor neurons in sham stimulated and CES stimulated mice. **c)** Longitudinal distribution of motor neurons relative to SCI site projecting to nerve transfer (right) or contralateral intact median nerve (left). Scale bar = 1 mm. Data presented as mean ± sem.

### CES enhances functional reinnervation of median nerve targets after nerve transfer

Acute electromyography (EMG) performed after 16 weeks of rehabilitation demonstrated that CES resulted in enhanced functional connectivity of musculocutaneous nerve motor axons to *FCR*. *FCR* responses showed lower thresholds for motor evoked potentials (MEPs) in the nerve transfer side of animals treated with CES (two-tailed t-test, p-value = 0.0408, t *=* 2.525, *df =* 8) (Figure 7). Suprathreshold stimulation resulted in similar MEP latency and amplitudes between CES and sham stimulation.

**Figure 7.**
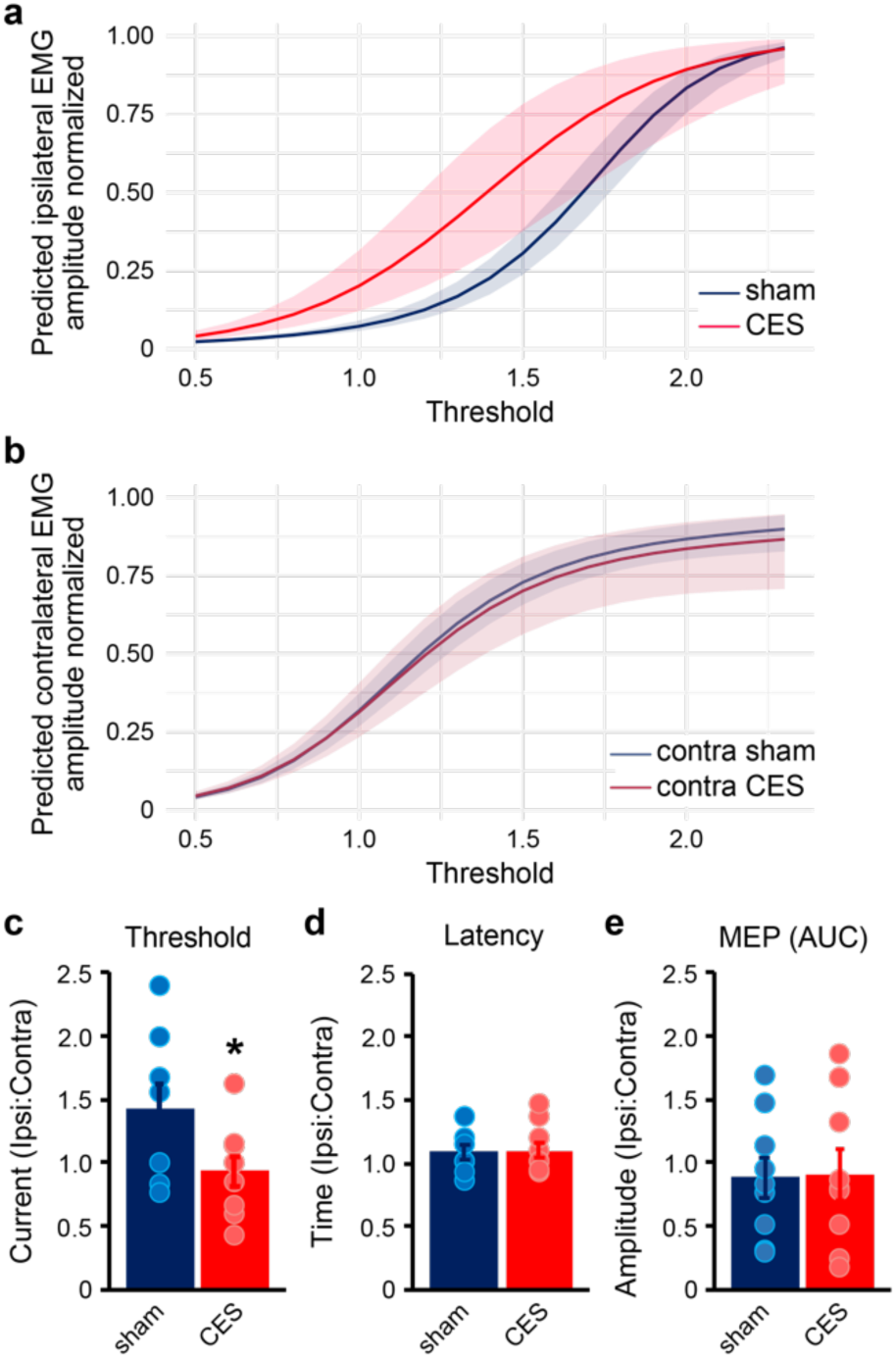
CES enhances functional reinnervation of median nerve target *FCR*. **a,b)** Predicted EMG response curves in *FCR* from stimulation of CES transfer nerves are similar to responses from stimulation of intact, contralateral median nerves. **c)** Evoked motor thresholds in CES transfer nerves normalized to intact, contralateral median nerve are lower than in sham stimulated mice (two-tailed t-test, *P* = 0.0408). **d,e)** Suprathreshold stimulation results in similar latency and MEP amplitude responses in sham or CES stimulated nerve transfers. Data presented as mean ± sem.

## Discussion

With a bedside to bench strategy, we generated a nerve transfer model to evaluate CES as a therapeutic approach to enhance outcomes in surgical intervention for chronic tetraplegia. Using a holistic HMFA, we found that CES altered the course of recovery for mice after nerve transfer surgery. We found both enhanced functional regeneration in response to CES, as well as considerable limits of the mouse as a model for nerve transfer to treat chronic SCI; however, within the context of the mouse model, we believe that our findings provide a strong rationale for advancing CES to a more clinically relevant model.

We observed a significantly enhanced rate of recovery on forelimb behavioral tests of naturalistic movements. CES treated mice showed more rapid recovery of pasta handling and tape removal. Following SCI, animals appeared to be able to use adaptive strategies to perform these behaviors without musculocutaneous innervation of forelimb flexors. In contrast, grip strength that depends on robust forelimb flexor activation was severely and permanently disrupted by nerve transfer, as expected. Additionally, the ballistic forelimb prehension movements of the single pellet reach task were never recovered after SCI. This likely arises from the absence of forelimb flexor activation during the reach. Without coordinated activation of the upper and lower forelimb, mice were unable to effectively execute the precisely coordinated movements required for the reaching task.

While behavioral recovery in sham stimulated animals eventually reached similar levels as in CES mice, there was still evidence of more effective reinnervation in CES mice after 16 weeks of rehabilitation. EMG thresholds were significantly lower in CES mice than sham controls. Combined with histological evidence of greater motor endplate innervation, this indicates more robust innervation of the median nerve target *FCR* in CES mice.

The size of the mouse nerves limited the granularity of peripheral circuit rewiring that was possible. In mice this meant sacrificing the full musculocutaneous nerve (motor and sensory) and targeting the entirety of the median nerve (pronation, wrist flexion, finger flexion, hand sensation, hand intrinsics), while in humans the brachialis branch of the musculocutaneous nerve can be used to specifically target finger flexion of the first 3 digits alone using the anterior interosseous branch of the median nerve alone (Brown, 2011). Size also tempered the pro-regenerative effects of CES relative to sham control. As the regeneration distances in mice are fractions of those necessary for effective regeneration in humans, the early behavioral improvements that we observed after CES were soon matched as axon regeneration caught up in sham animals.

These shortcomings do not indicate a failure of the strategy, but rather provide us with a more thorough understanding of the underlying circuit mechanisms that shape outcomes from nerve transfer. By integrating knowledge gained from our animal studies with clinical observations and participant feedback in our clinical research, this bidirectional approach enables a more comprehensive understanding of the strategies required for improved outcomes for individuals living with tetraplegia.

## Data Availability statement

Full dataset generated during this study will be available upon acceptance for publication in a publicly accessible G-Node GIN repository found at https://gin.g-node.org/Circuit_Repair_Lab and the Open Data Commons for Spinal Cord Injury (ODC-SCI, lab ID 110, Hollis lab).

## Acknowledgements

This work was supported by the BNI Structural and Functional Imaging Core, Paralyzed Veterans of America Research Foundation PVA22_R_00037, the New York State Department of Health Spinal Cord Injury Research Board C34463GG, and the National Institutes of Health DP2 NS106663, S10 OD028547.

## Author contributions

J.S.J., J.B., and E.H. conceived and designed the project; J.S.J., M.S., A.B., and P.diG. performed the experiments and collected data; A.R.F and A.T-E. conceived and designed analytical pipeline; J.S.J, M.S., A.B., and A.T-E. performed data analysis and interpreted results; J.S.J., M.S., A.B., A.T-E., and E.H. wrote the manuscript.

## Notes

### Competing Interest Statement

The authors have declared no competing interest.

